# Signal transfer of visual stimuli to V4 occurs in gamma-rhythmic, pulsed information packages

**DOI:** 10.1101/2020.02.03.931956

**Authors:** Dmitriy Lisitsyn, Iris Grothe, Andreas K. Kreiter, Udo A. Ernst

## Abstract

**Summary:** Selective visual attention allows the brain to focus on behaviorally relevant information while ignoring irrelevant signals. As a possible mechanism, routing by synchronization was proposed: neural populations sending attended signals align their gamma-rhythmic activities with receiving populations, such that spikes from the senders arrive at excitability peaks of the receivers, enhancing signal transfer. Conversely, the non-attended signals arrive unaligned to the receiver’s oscillation, reducing signal transfer. Therefore, visual signals should be transferred through periodically pulsed information packages, resulting in a modulation of the stimulus content within the receiver’s activity by its gamma phase and amplitude. To test this prediction, we quantified gamma phase-specific stimulus content within neural activity from area V4 of macaques performing a visual attention task. For the attended stimulus we find enhanced stimulus content reaching its maximum near excitability peaks, with effect magnitude increasing with oscillation amplitude, establishing a functional link between selective processing and gamma activity.

## Introduction

Visual information processing is computationally demanding, requiring the brain to handle a continuous, high-dimensional stream of sensory input signals. Selective attention helps to reduce this computational complexity by focusing on signals which are behaviorally relevant at the expense of other, currently irrelevant signals.

Evidence for such selective processing is already apparent at the single neuron level: when presented with two stimuli inside their receptive fields (RFs) of which only one is behaviorally relevant and attended, V4 neurons respond as if only the attended stimulus was present (Moran and Desimone, 1985; Reynolds et al., 1999). At the same time, there is only a small attention-dependent modulation of the firing rates of the populations in V1/V2 providing the input signals from the two visual stimuli to V4 (Luck et al., 1997; McAdams and Maunsell, 1999; Mehta et al., 2000; Moran and Desimone, 1985; Motter, 1993; Salinas and Sejnowski, 2000). This suggests that selective responses in downstream areas do not result from strong upstream rate modulations, like silencing the population of afferent neurons conveying distractor-related signals. Hence, selective attention in downstream areas must rely on a different mechanism for dynamically changing effective connectivity depending on task demands. One influential idea to explain such effects relies on inter-areal synchronization of oscillatory neural activity in the gamma-band (30Hz-100Hz) in order to enact dynamic modulation of effective connectivity between the V1 and V4 neurons (Fries, 2015, 2005; Kreiter, 2006), with proof of concept provided by a range of modeling studies (e.g. Akam and Kullmann, 2010; Börgers and Kopell, 2008; Buehlmann and Deco, 2010; Harnack et al., 2015; Palmigiano et al., 2017).

Synchronization in the activity of a presynaptic (sending) population increases its post-synaptic influence by making the neurons’ spiking outputs coincident in time, an effect found to be enhanced for populations processing attended versus non-attended stimuli (Azouz and Gray, 2003; Bichot et al., 2005; Fries et al., 2001; Steinmetz et al., 2000; Taylor et al., 2005). For the post-synaptic receiver population, local oscillatory synchronization establishes a rhythmic modulation of gain, with alternating periods of high and low excitability (Atallah and Scanziani, 2009; Buzsáki and Wang, 2012; Ni et al., 2016; Salkoff et al., 2015; Vinck et al., 2013). Noteworthy, the amplitude of gamma oscillations is not constant, for instance showing modulation by a low-frequency rhythm in the 6-10 Hz range (Bosman et al., 2009; Fries, 2015; McLelland and VanRullen, 2016; Palmigiano et al., 2017; Spyropoulos et al., 2018). It is reasonable to expect that the gain modulation by gamma phase is then itself higher during periods of high-amplitude oscillatory activity and lower during periods of low-amplitude oscillatory activity.

If both sending and receiving populations exhibit oscillatory activity, they can establish coherent states in which their phases are coupled (see insets in Fig. 1A). In a favorable state for information routing, bursts of spikes from a presynaptic subpopulation arrive at the receiving population close to its peak of excitability, resulting in enhanced information transfer (left inset of Fig. 1A). Conversely, in an unfavorable phase relationship, the sending population’s bursts arrive predominantly during the windows of inhibition in the receiver population, and information transfer is suppressed (first three cycles in the right inset of Fig. 1A). In consequence, effective connectivity can be modulated by establishing the appropriate phase relationship between the rhythmic activities of the presynaptic (sending) and post-synaptic (receiving) populations. This mechanism is in general being referred to as communication-through-coherence (CTC) (Fries, 2015, 2005), or as routing-by-synchrony (RBS) (Grothe et al., 2018, 2012; Kreiter, 2006; Palmigiano et al., 2017). Two studies demonstrating such a dynamic routing mechanism showed that V4 populations do indeed establish a stronger phase coherence with the presynaptic populations processing the attended stimulus than those processing the non-attended stimuli (Bosman et al., 2012; Grothe et al., 2012). Further evidence in support for CTC and RBS has also been reported across other visual areas (Besserve et al., 2015; Jia et al., 2013; Womelsdorf et al., 2007) and different brain regions (Buschman and Miller, 2007; Cardin et al., 2009; Siegle et al., 2014).

**Figure 1.**
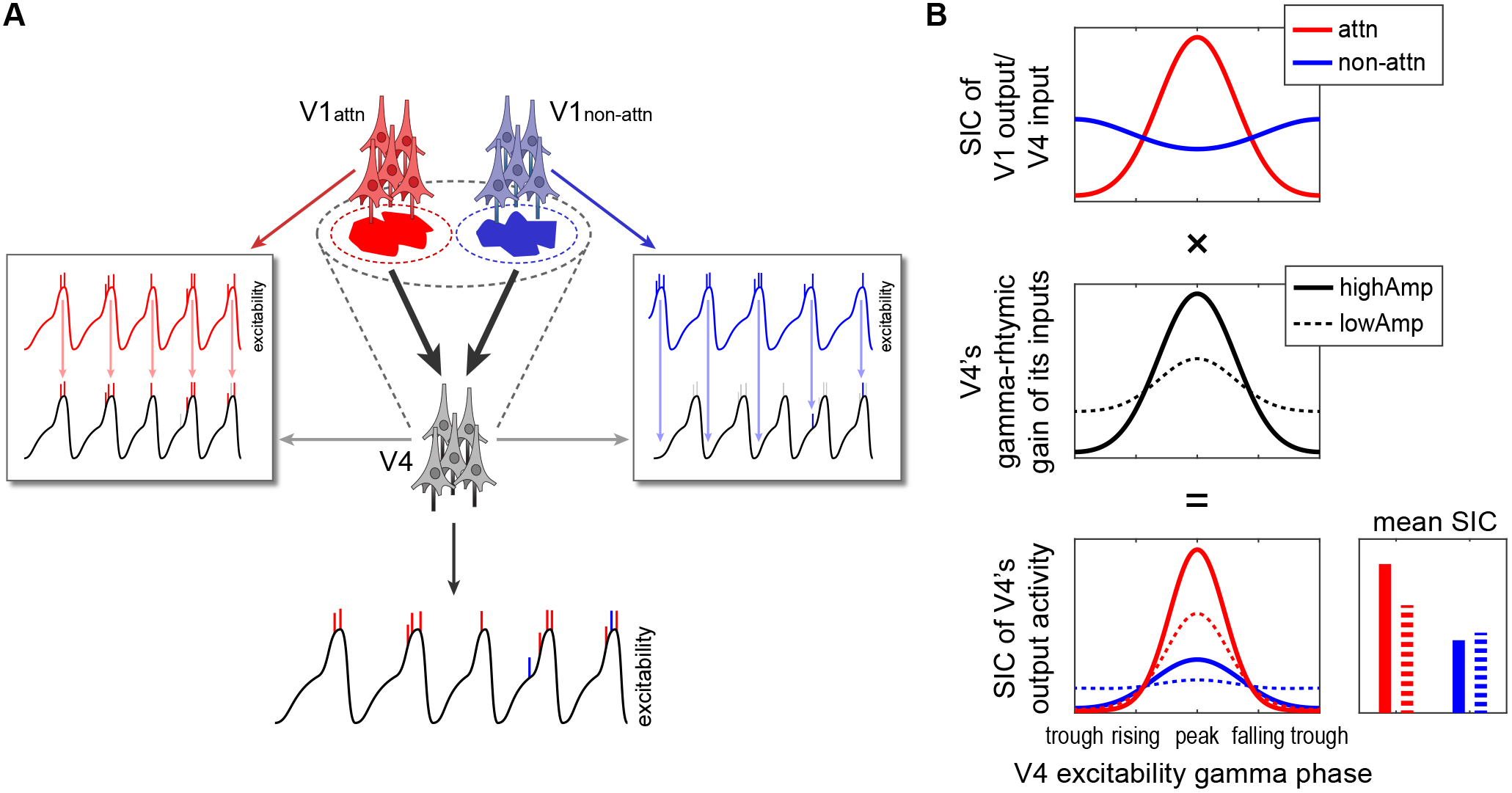
Routing by synchrony mechanism predicts modulation of stimulus information content within V4’s gamma-rhythmic neural activity, depending on V4’s gamma phase and amplitude. (A) Schematic outline of the routing by synchrony (RBS) mechanism, which allows downstream neurons to process stimulus information mediated by selected subsets of afferent inputs while suppressing the information provided by other inputs. Two stimuli (red and blue shapes) compete for being processed by a downstream V4 population with a large receptive field (RF) containing both stimuli indicated by the gray dashed oval. Each visual stimulus is contained within RFs of separate V1 populations (red and blue dashed ovals) evoking spiking activity within their corresponding V1 population (red and blue vertical bars in the insets). The V1 populations exhibit gamma-rhythmic activity, causing their spikes to occur in bursts. These bursts of spikes act as input to V4, which exhibits its own gamma-rhythmic activity (gray oscillatory lines in the insets). The rhythmic activity of the V1 population corresponding to the attended stimulus is synchronized with V4’s gamma rhythm in a favorable phase relationship, such that its spikes arrive at V4 when it is most excitable (left inset, red arrows). This effectively evokes spiking activity in V4, resulting in a reliable transfer of the attended stimuli’s information. Conversely, the rhythmic activity of the V1 population with the non-attended stimulus in its RF exhibits substantially less phase-locking with V4’s gamma rhythm (Bosman et al., 2012; Grothe et al., 2012), resulting in many cycles where the bursts of spikes arrive at V4 when it is least excitable, failing to evoke further spikes (right inset, light blue arrows). In consequence, the transfer of the non-attended stimulus signal is suppressed. (B) Scheme showcasing how the information contained within attended and non-attended stimuli (stimulus information content, SIC), should be modulated depending on V4’s gamma phase and amplitude in accordance with RBS. Assuming that the upstream cortical population processing the attended stimulus establishes a favorable phase relationship with V4 as shown in (A, left inset), the highest amount of attended SIC should arrive during V4’s excitability peak (red line in top plot). Assuming that for the non-attended stimulus, the corresponding upstream population establishes substantially less phase-locking with V4, and in a predominantly anti-phasic relationship, we expect SIC modulation to be much lower, with a slightly higher amount of non-attended SIC arriving at V4’s excitability trough than at its peak (blue line in top plot). The middle plot shows V4’s gamma-rhythmic gain (solid black line for high amplitude gamma activity and dashed black line for low amplitude). Modulating the phase-specific SIC inputs to V4 (top plot) by V4’s gain (middle plot) provides a prediction of how attended and non-attended SIC should depend on V4’s excitability gamma phase and amplitude within its output activity (bottom plot). The corresponding bar plot (bottom right) displays the average SIC within V4’s output activity independent of phase, demonstrating how attended and non-attended SIC should change during V4’s high amplitude versus low amplitude gamma activity (solid colored bars for high amplitude condition and dashed colored bars for low amplitude condition).

As a direct implication, RBS posits that stimulus information arrives at V4 from lower visual areas in the form of *periodically pulsed information packages*, where the phase relationship between the rhythmic activities of the sending and receiving populations determines whether the packages get passed on or will be suppressed. Behavioral correlates compatible with such a pulsed information transfer were reported by two recent studies demonstrating that reaction times to a sudden stimulus change depend on both, the V4 gamma phase just before the transient response evoked in V4 by the stimulus change (Ni et al., 2016), and on the V1-V4 phase relationship preceding the stimulus change (Rohenkohl et al., 2018). These results are line with the assumption that transient input arriving during V4 excitability peaks could improve behavioral responses.

Here, we set out to directly show that selective information transfer of extended, time-varying signals relies on series of packets that enter processing in V4 depending on excitability phase and amplitude. For this purpose the stimulus paradigm introduced by Grothe et al., 2018 provides a unique opportunity to experimentally test the predictions of a ‘pulsed’ routing scheme and to scrutinize the RBS mechanism. In the experiment, two stimuli, attended and non-attended, presented within the RF of a local neural population in V4 were tagged with independently varying luminance signals, allowing to measure the stimulus information content (SIC) within V4’s activity concurrently for both input signals over prolonged periods of time (Fig. 2). Quantifying SIC by a frequency-resolved correlation measure (spectral coherence) between input signals and recorded neural activity allowed to show that SIC for the attended stimulus was strongly enhanced relative to the SIC for the non-attended stimulus.

In the present study, we utilize this data set in order to quantify the SIC of neural activity occurring at different phases of successive gamma-oscillation cycles. This allows us to quantify how well signals of attended and non-attended stimuli are routed to downstream populations during different phases of the V4 gamma-oscillation cycles at which they arrive and how this depends on oscillation amplitude. If gamma synchronization and selective information routing were causally linked to each other according to RBS, the SIC for attended and non-attended stimuli should be modulated by phase and amplitude of V4’s gamma-rhythmic activity (Fig. 1B). In particular, for the attended stimulus, we expect SIC to be modulated by phase such that the highest levels correspond to V4’s excitability peaks. This SIC modulation by phase should decrease during periods of low gamma activity. For the non-attended stimulus, the modulation strength should be consistently lower than that of the attended stimulus. Further, the largest differences between attended and non-attended SIC should occur at the excitability peak during high amplitude oscillatory activity.

**Figure 2.**
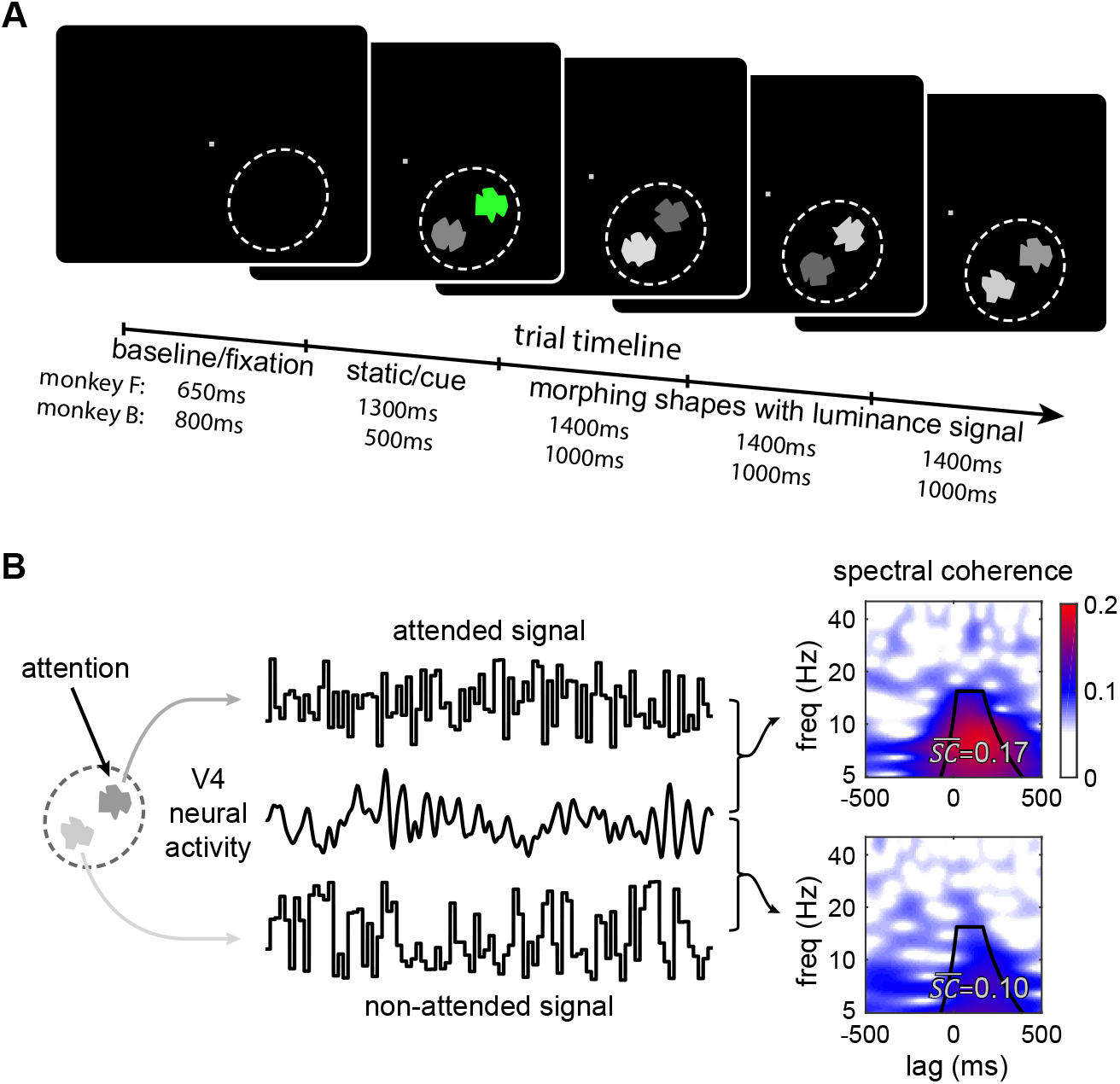
Behavioral task and quantification of stimulus information content. (A) Stimulus sequence. After the monkey presses a lever, the trial starts with the appearance of a fixation spot (baseline period). Shortly afterwards, two stimuli in the form of static shapes are presented within the RF of the V4 recording site (dashed ellipse). One of the shapes is cued to be memorized and attended with green shading while the other shape has to be ignored (static period). Then the cued shape reverts to gray and both stimuli begin to morph into different shapes. After a number of morph cycles, the initially cued shape reappears in the attended location. If the animal releases the bar within a short time window around the reappearance of the cued shape, a reward is delivered. (B) Throughout the morphing period, each shapes’ luminance was modulated in time by a random white-noise signal. These luminance fluctuations were irrelevant to the task but served as independent tags for signals originating from the stimulus to be attended, and from the stimulus to be ignored. We evaluated spectral coherence between the recorded neural signals and each input signal to quantify stimulus information content (SIC) in V4 activity. By pooling across relevant lag and frequency bins from the spectral coherence (indicated by the black lines) a single value (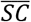) was acquired for each signal.

## Results

For investigating whether and how attention-dependent signal transfer is affected by V4’s gamma phase and amplitude, we analyzed local field potentials (LFP) and multi-unit spiking activity (measured as entire spiking activity, ESA) recorded from the superficial layers of area V4 in two macaque monkeys (Macaca mulatta). During recording, the animals were engaged in a demanding shape tracking task requiring the monkeys to attend to one of two concurrently presented dynamic stimuli within the recorded population’s receptive field (RF) (see Fig. 2A and methods section for details). The two stimuli consisted of complex shapes, which, after an initial static period, morphed through a series of different shapes throughout the trial. At the beginning of each trial, one of the two stimuli was cued. The task for the monkey was to attend to the cued stimulus while maintaining fixation, and to respond when its initial shape reappeared in the morphing sequence. The other, non-attended stimulus had to be ignored. The number of morph-cycles that the stimuli went through before returning to the initial shape was randomized.

Crucially, the two stimuli were tagged by independent and behaviorally irrelevant random luminance fluctuations, with a luminance change every 10ms. This allows us to evaluate stimulus information content (SIC) in V4 activity by computing the spectral coherence between the neural activity and the luminance signals (see Fig. 2B). Spectral coherence provides a frequency- and time-delay-resolved correlation measure between two signals. By pooling across relevant lag and frequency bins, we acquired a single value 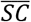 as a measure for the average SIC of the shapes’ luminance fluctuation within the recorded neural activity. The neural signals included in this analysis were taken from the start of the second morph cycle until 200 ms before a correct behavioral response. The first cycle was excluded since it never morphed into the target shape and thus would not require the animal to pay ‘full’ attention to the target shape in this particular time interval.

In order to probe whether SIC is modulated by phase and amplitude as predicted in figure 1B, we employed ESA as a proxy for excitability. Specifically, we first identified which LFP gamma phase was associated with maximum ESA, and then shifted the LFP gamma phase by the appropriate amount for each recording site giving us the ESA-aligned gamma phase. Throughout the whole analysis, this is the gamma phase employed as a proxy for excitability’s gamma phase (Fig. 3A).

**Figure 3.**
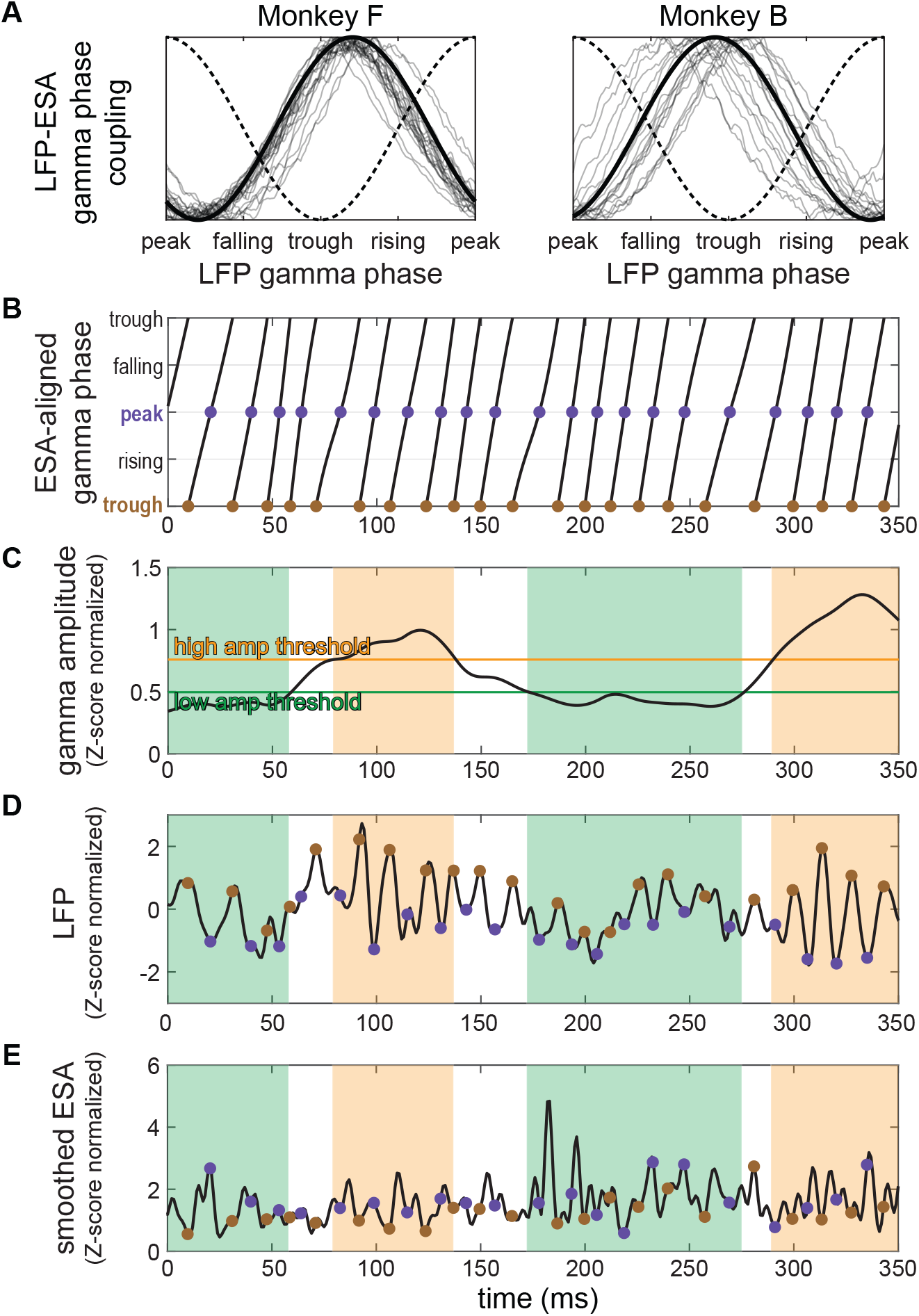
Extraction of gamma phase-and amplitude-specific neural signals. (A) Normalized spike-field coupling in the gamma frequency range, computed from ESA and LFP signals, for all the recording sites for each animal (thin lines for each site, bold line for the mean across sites). The peak of spiking activity consistently occurs at roughly the trough of LFP’s gamma cycle (indicated by dotted line), as expected for the superficial layer (van Kerkoerle et al., 2014). By shifting LFP’s gamma phase by the appropriate amount, individually for each session, we attain the ESA-aligned gamma phase, which is used as a proxy for excitability gamma phase throughout the rest of the analysis. (B) Time course of ESA-aligned gamma phase corresponding to a 350ms snippet of a trial. The locations of peaks are marked with purple dots, and excitability troughs with brown dots. (C) Corresponding gamma amplitude with 30% highest and 30% lowest thresholds marked with horizontal lines. Periods of time when the amplitude surpassed the high threshold are shaded in orange, and periods of time below the low amplitude threshold are shaded in green. (D) The corresponding LFP neural activity with precise peak and trough times of the ESA-aligned gamma phase identified in (B), and high and low amplitudes identified in (C). By using the corresponding samples of the neural activity (either purple or brown dots for peak vs trough, or orange or green time periods for high vs low amplitude, or a combination of both), we can compute the amount of SIC within the selected components of the neural activity. Note that the LFP is obtained by low-pass filtering the recorded signal and thus each value represents neural activity from a small time window around it. (E) Same as in (D), but for the corresponding ESA signal

To quantify SIC in dependence on gamma phase and amplitude, we extracted neural activity specific to each phase of the gamma cycle and separately for periods of high and low gamma-band amplitudes. We then computed 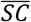 between these phase and amplitude-specific signals and the luminance fluctuations of the attended and non-attended stimuli (see Fig. 3 and methods section for details).

### Gamma phase modulates signal information content of the attended stimulus

In order to estimate how SIC depends on the gamma phase of V4, we extracted components of the neural signals associated with a specific gamma phase, by selecting the discrete time points that correspond to that particular phase, and sampling the neural signals at those points. In the example shown in figure 3B, we marked the time points corresponding to ESA-aligned gamma peaks (in purple) and troughs (in brown). The dots in figures 3D and E indicate which samples from the LFP and ESA signals, respectively, will be obtained when selecting at peaks (in purple) or troughs (in brown). The method is not limited to sampling just from the peak or from the trough, allowing to extract a signal specific to any desired phase. By computing 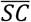 between the phase-specific neural activity signals and the input stimuli, we can assess the amount of attended or non-attended SIC in dependence on gamma phase.

For the attended stimulus (Fig. 4A, left two columns), SIC at peaks (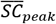) was significantly larger than SIC at troughs (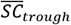) (p<0.00005 for all conditions, non-parametric permutation test). This result was consistent across monkeys and signals (LFP and ESA). In contrast, for the non-attended stimulus, there was no significant difference between SIC at peaks and troughs for ESA. The LFP showed a significantly higher SIC at peak vs trough (p=0.01 for monkey F LFP, p=0.0005 for monkey B LFP), but with a much smaller modulation than observed for the attended stimulus. Aside for a few values near the ESA-aligned gamma trough for monkey B’s non-attended ESA results, all observed 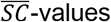 were significantly greater than chance level, showing that information is usually transferred during all gamma phases and never completely blocked.

**Figure 4.**
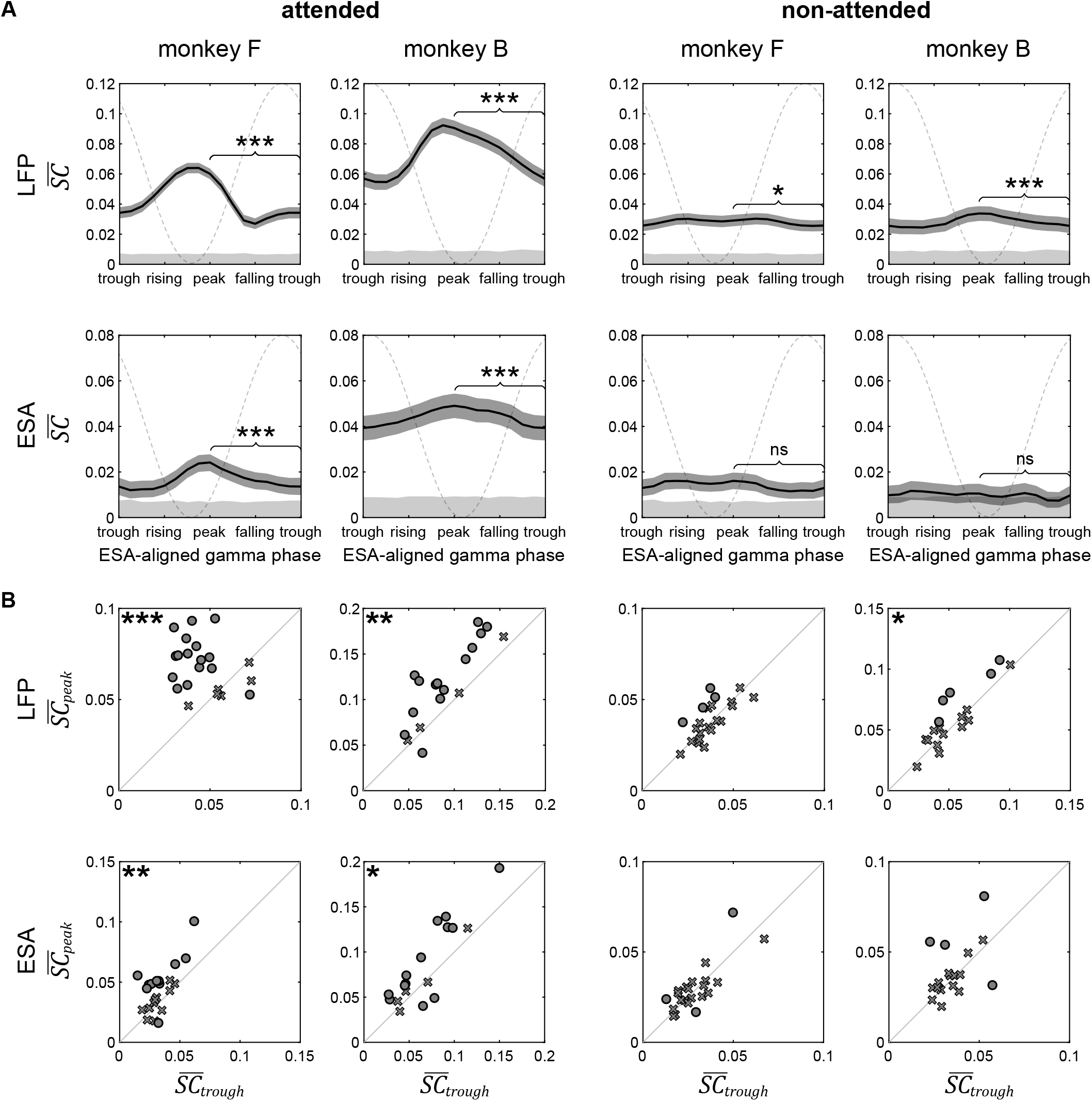
Stimulus information content is modulated by V4’s gamma phase. (A) SIC dependence on phase for data pooled across all sessions. In each plot, we display how 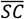 depends on the ESA-aligned gamma phase (horizontal axis) from which the neural signal is extracted. The shading around each line corresponds to the 95% confidence interval. Significance of the difference between 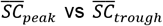 is indicated in each plot (*p<0.05, **p<0.01, ***p<0.001). The gray shading at the bottom of each plot corresponds to the 95% chance level (values below this level indicate no significant SIC). The grey dashed sinusoid line indicates the corresponding average LFP phase. (B) SIC at peaks versus troughs of the ESA-aligned gamma phase for individual sets. For each condition, we display a scatter plot of 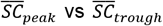 pairs. In the scatter plots, individual sets that exhibit a significant difference (p<0.05) are marked with a circle, and with a cross otherwise. The significance of the group distribution, i.e. whether it lies significantly below or above the diagonal, is marked with black asterisks on the side that contains significantly more trial sets (*p<0.05, **p<0.01, ***p<0.001).

The effect of a higher SIC at peaks than at troughs for the attended stimulus was consistent across individual recording trial sets. The scatter plots in the two leftmost columns of Fig. 4B display 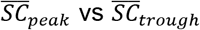 for each individual trial set for the attended stimulus. The majority of data points show significantly higher SIC at peaks rather than troughs (p<0.05, marked with circles; 15 of 22 for monkey F LFP, 13 of 18 for monkey B LFP, 10 of 22 for monkey F ESA, and 11 of 18 for monkey B ESA). Note that only a few trial sets exhibit significantly lower SIC at gamma peaks (1 of 22 for monkey F LFP, 1 of 18 for monkey B LFP, 1 of 22 for monkey F ESA, 2 of 18 for monkey B ESA).

We assessed the individual set’s group statistics by determining whether the distribution of log-ratios, 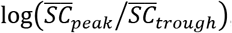, is significantly higher or lower than 0, indicated by the black asterisks in the scatter plots of Fig. 4B (Wilcoxon signed rank test). For the attended stimulus, the peak SIC was significantly higher than the trough SIC across both monkeys and both neural signals (p=0.0003 for monkey F LFP, p=0. 0013 for monkey B LFP, p=0.0097 for monkey F ESA, and p=0.012 for monkey B ESA). For the non-attended stimulus, the scatter plots show a tendency to exhibit higher 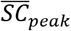 than 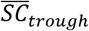 with more significant results above the diagonal, however, the group statistic was only significant for monkey B’s LFP (p=0.041, top right scatter plot in figure 4B).

### Gamma amplitude modulates signal information content of the attended stimulus

For the amplitude dissection, we first gathered the distribution of gamma amplitudes throughout each individual trial set. From this distribution, we selected the 70th (and 30th) percentiles to use as high (and low) cutoff thresholds to select activity from high and low gamma amplitude periods. Using these thresholds, for each trial, we selected the time periods exhibiting high (or low) oscillation amplitudes. In the examples in figure 3C, the corresponding periods are indicated by orange and green shading, respectively. In figures 3D and E, the same shading indicates which periods of the LFP and ESA signals were selected by amplitude dissociation. We next evaluated and compared the amount of SIC within high amplitude (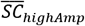) versus low amplitude gamma oscillations (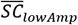).

When we ran the analysis on a cumulative set of all the trials over all recording sites for the attended stimulus, we found a small but significant increase of SIC in neural activity during high amplitude gamma oscillations in comparison to low amplitude gamma activity (Fig. 5A). Again, this result holds for all monkeys and neural signal types (p=0.011 for monkey F LFP, p<0.00005 for monkey B LFP, p=0.0003 for monkey F ESA, p=0.0001 for monkey B ESA). This corresponds to the prediction derived from our hypothesis (see Fig. 1B). For the non-attended stimulus, the analysis did not reveal any significant differences between SIC within periods of high versus low gamma activity.

**Figure 5.**
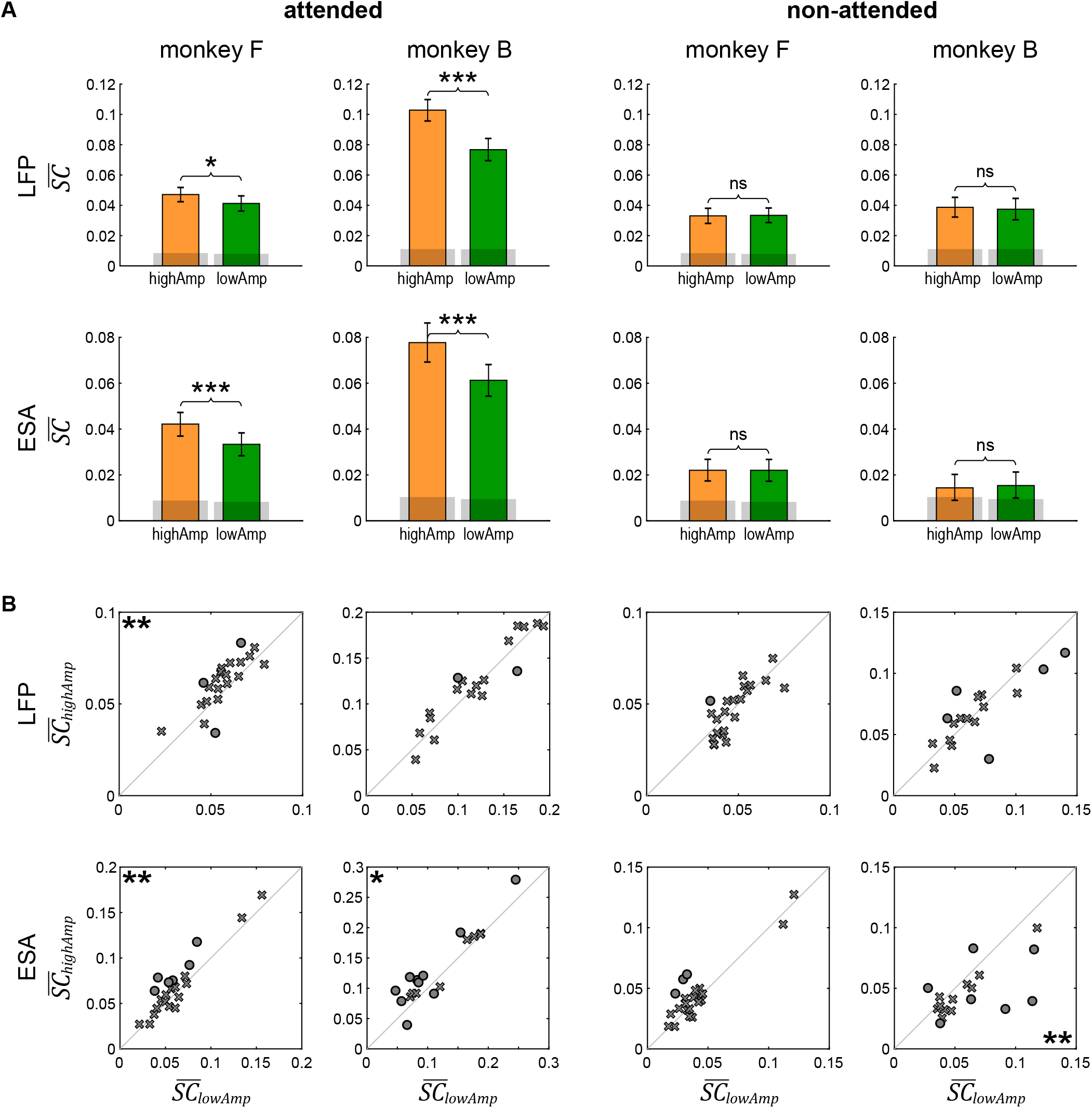
Stimulus information content is modulated by V4’s gamma amplitude. (A) SIC extracted from periods with high- versus low-amplitude gamma oscillations for data pooled across all recording sessions. In each plot, for the specific condition as indicated by the row and column labels, we display pairs of 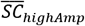 (in orange) and 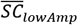 (in green), separately for each animal, neural data type, and attention condition. The error bars indicate the 95% confidence interval. The gray shading at the bottom of each bar indicates the 95% chance level for that value. Significance level of the differences is indicated above each pair of bars, computed via permutation testing across the trials (*p<0.05, **p<0.01, ***p<0.001). (B) SIC extracted from periods of high versus low gamma neural activity for individual sets. For each condition, we display a scatter plot of 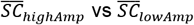 pairs. In the scatter plots, sets that exhibit a significant difference (p<0.05) are marked with a circle, and with a cross otherwise. The significance of the group distribution, whether it lies significantly below or above the diagonal, is marked with black asterisks on the side that contains significantly more sets (*p<0.05, **p<0.01, ***p<0.001).

Results from analyzing individual sites corroborated the cumulative outcomes (Fig. 5B). For the attended stimulus, the majority of the sites with a significant difference did indicate increased information present during periods of high amplitude oscillations as seen in the scatter plots in the left two columns of Fig. 5. When we looked at the distribution of the ratios 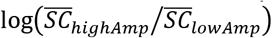, except for LFPs from monkey B, all conditions exhibited a significant shift towards a higher SIC for high amplitude gamma, as indicated by the black asterisks (p=0.0047 for monkey F LFP, p=0.0089 for monkey F ESA, p=0.011 for monkey B ESA).

For the non-attended stimulus (right two columns of Fig. 5), SIC did not show any significant differences for the individual sets, except for monkey B’s ESA signal. This was the only case for which we found that 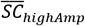 is significantly *lower* than 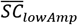 (p=0.0083).

### Stimulus information content modulation by phase is increased during higher gamma amplitude activity

Dissections with respect to phase and amplitude were combined to select LFP and ESA activity associated to the co-occurrence of a particular gamma phase and amplitude (Fig. 3).

In figure 6, the results for the SIC computed from the set of all trials collected from all the recording sites is displayed, with the high amplitude results in orange and low amplitude results in green. Corroborating the hypotheses in Fig. 1B, the data exhibited higher SIC modulation by gamma phase within high amplitude periods and lower SIC modulation by phase for the low amplitude periods.

**Figure 6.**
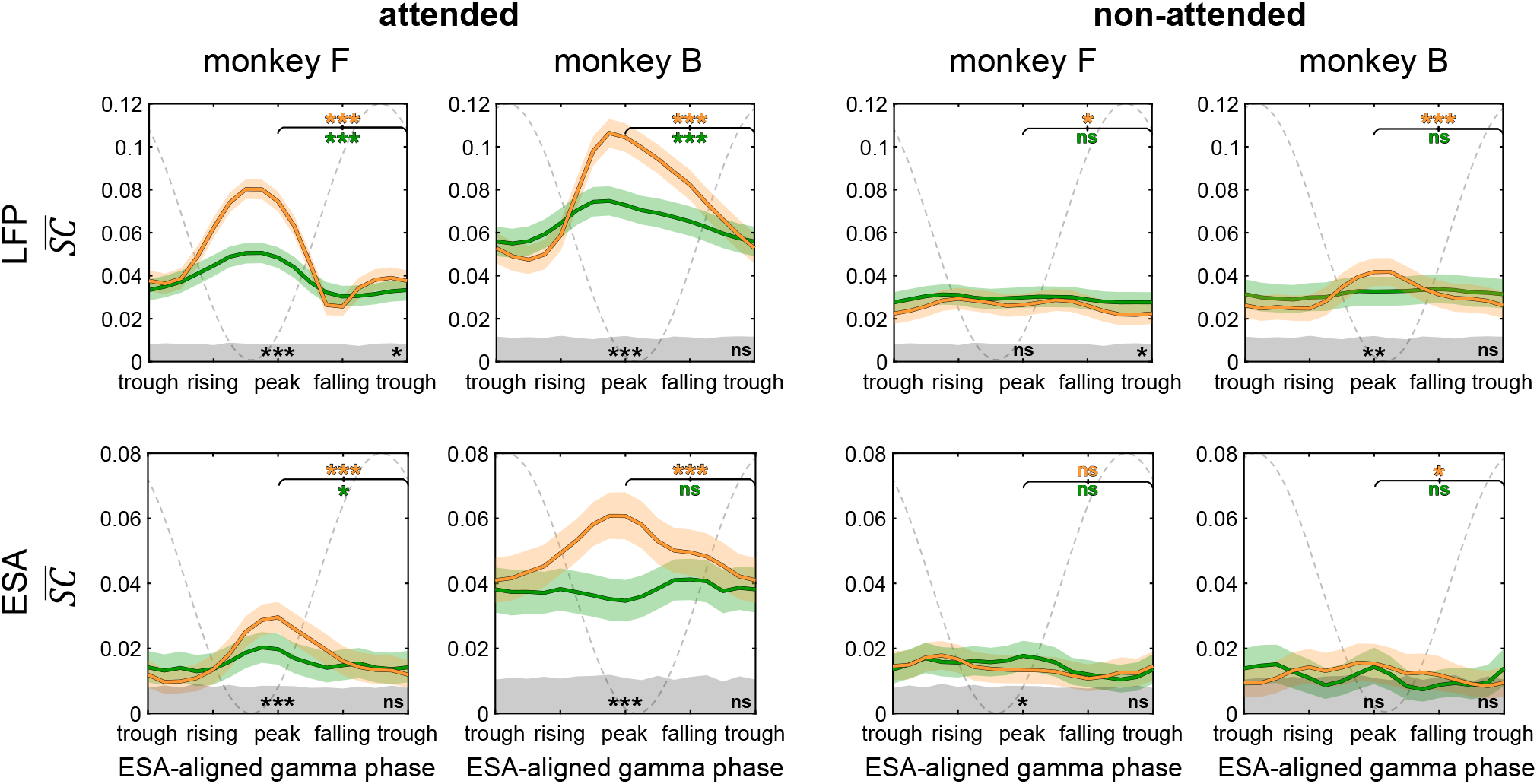
Signal information content during high and low gamma amplitude periods in dependence on V4’s gamma phase. For each condition, as indicated by the row and column labels, each plot displays how SIC is modulated by phase from which the neural signal is extracted from (horizontal axis), separately for high gamma amplitudes (in orange) and low gamma amplitudes (in green) conditions. The shading around each line corresponds to the 95% confidence interval. The grey shading at the bottom of each plot shows the corresponding 95% chance level. The grey dashed sinusoid line indicates the corresponding average LFP phase. The asterisks towards the bottom of each plot indicate whether there is a significant difference between 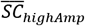 and 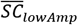 for the peak or the trough. The colored asterisks at the top right of each plot indicated whether there is a significant difference between 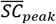 and 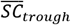 for the corresponding amplitude condition.

The outcome lends itself to multiple comparisons: comparing SIC at high versus low amplitudes at peak and trough (significance indicated at the bottom of each plot at peak and trough phases in black) and comparing SIC at peak versus trough phases in either high or low amplitude conditions (significance indicated at the top of the plot with colors corresponding to the amplitude conditions).

The difference between high and low amplitudes for the attended stimulus was strongly phase-specific: while SIC near peaks within high gamma amplitude activity was significantly larger than SIC within low gamma amplitude activity (p<0.00005 for both animals and both neural signals), there were no significant differences near troughs except a small one for monkey F’s LFP result (p=0.045). Consequently, SIC modulation by gamma phase was higher in the high amplitude condition (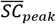 is greater than 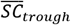 with p<0.00005 for all data sets), and lower (but still significant in three out of four cases) in the low amplitude condition (p<0.00005 for monkey F and monkey B LFP, p=0.012 for monkey F ESA and no significant different for monkey B LFP). Overall, this corroborates the prediction from the corresponding curves in figure 1B: for the attended signal, SIC modulation by phase is increased during high amplitude gamma activity and decreased during low amplitude gamma activity.

Similar to the previous phase-only analysis (see Fig. 4), SIC modulation by phase was reduced dramatically for the non-attended stimulus when compared to the attended stimulus (Fig. 6 right two columns). However, in the high amplitude condition, we observed more cases with a significantly higher SIC at the peak versus the trough (p=0.019 for monkey F LFP, p<0.00005 for monkey B LFP, p=0.016 for monkey B ESA). In the low amplitude condition, the differences between peaks and troughs became insignificant. Simultaneously, we observed for a few particular phases that the SIC is higher in the low amplitude condition than in the high amplitude condition (at the troughs for monkey F LFP with p=0.033 and at the peaks for monkey F ESA with p=0.017).

## Discussion

In the present work we tested the prediction of the RBS mechanism that attention-dependent visual stimulus information is not continuously dispersed over time but instead occurs selectively in pulsed packages, modulated by phase and amplitude of V4’s gamma-rhythmic activity.

Our analysis reveals that the luminance fluctuations tagging the attended stimulus are expressed most strongly within neural activity close to the phase in the gamma oscillation cycle where local spiking peaks (or close to the LFP troughs). During periods with high amplitude gamma oscillations the overall SIC for attended stimuli is higher in the V4 activity than during low amplitude oscillations, whereas for non-attended stimuli there is no significant difference. SIC’s increase with amplitude is particularly strong at the spiking activity peak and absent at the trough. Since the peaks of spiking activity should roughly correspond to the peaks of excitability, these results corroborate central predictions derived from the RBS mechanism.

Certain details of the results seem to deviate from the predictions illustrated in figure 1B. For instance, when comparing attended and non-attended SIC, our expectation was that the non-attended SIC should be at least as large as the attended SIC in the vicinity of excitability troughs. However, aside for monkey F’s ESA result, we find that the attended signal is consistently better expressed across almost all phases. This may be explained by a temporal dispersion of information across phase, which occurs due to multiple factors. First, the precision of phase estimation itself is limited by noise in the recorded neural activity. Additionally, we also smoothed the neural activity prior to the phase-specific signal extraction, such that the value of the signal at each time point represents a temporal window of activity around it. Finally, computing SIC based on spectral coherence involves comparing the neural recording and luminance fluctuation signals with wavelets that are centered at the time of interest but also extend in temporal space. Taken together, these methodological limitations all lead to the luminance flicker signal being partially mapped to phase ranges at which it does not actually occur, thereby increasing SIC in low-SIC ranges and decreasing SIC in the high-SIC ranges of the oscillatory cycle. In consequence, our analysis consistently underestimates the magnitude of SIC modulation by phase.

Dissociating signal content with respect to gamma amplitude could be subject to a similar reduction in SIC modulation amplitude. If we assume a constant noise level throughout the recordings, precision of phase estimation will decrease during periods of low gamma amplitude activity. Such an effect may contribute to the differences between high and low amplitude SIC results observed in figure 6 where we perform a simultaneous phase and amplitude-specific signal extraction.

The results for the non-attended signal provide further insight into the details of the RBS mechanism. Here, we observe only weak, or no phase-dependent modulation of SIC. The absence of a strong modulation implies a certain coherence for the non-attended signal with the receiving population in an anti-phasic relationship in order to counteract V4’s gain modulation. This is compatible with a weak level of coherence between non-attended V1 and V4 as observed in Grothe et al., 2012. On the other hand, a weak modulation of non-attended SIC in V4 could emerge if there is no coherence between non-attended V1 and V4 as observed in Bosman et al., 2012. In principle, our results are also compatible with a strong anti-phasic coherence between sender and receiver, however, such a strong coherence has never been observed experimentally, even though it would be functionally optimal for selective information routing. This may be due to how the phase-locked states between the sending and receiving populations are established. Previous studies have shown evidence that gamma activity acts in a feedforward manner, with the upstream populations’ gamma rhythm entraining the gamma in downstream V4 (Bastos et al., 2015; Bosman et al., 2012; Michalareas et al., 2016; Richter et al., 2017; van Kerkoerle et al., 2014). With this in mind, it is up to the upstream population processing the attended stimulus to entrain its downstream V4 with its own gamma rhythm, whereas the population processing the non-attended stimulus fails to entrain V4, remaining primarily uncoupled.

Contrary to the proposed RBS mechanism, an explanation for the occurrence of pulsed information packages may solely rest on the oscillatory activity of the receiver neurons. Having more spikes in the vicinity of V4 excitability peaks has the potential to encode more information, irrespective of any inter-areal phase coherence between sending and receiving population. However, the absence of a strong modulation for the non-attended stimulus in our data suggests that a trivial rate modulation in the sender populations cannot explain our results. Further, it is important to note that this potential rate-coding confound is not in any way in contradiction with the RBS mechanism. As long as the stimulus information is transmitted between populations by synchronizing the bursts of activity from the sender with the peaks of excitability of the receiver, whether or not these bursts exhibit rate-coding effects, the pulsed transmission scheme holds. Further, since rates and SIC in the sending populations are barely modulated by attention (Luck et al., 1997; McAdams and Maunsell, 1999; Mehta et al., 2000; Moran and Desimone, 1985; Motter, 1993), without selective phase coherence we would expect to observe very similar phase modulation strengths for both, attended and non-attended SIC.

A direct impact on behavior by the pulsed information transmission scheme of RBS might occur if the animals can not accumulate SIC over prolonged periods of sustained attention, as in our experimental setup. For example, neural responses to rapid stimulus changes exhibit fast transients lasting only few gamma cycles (Traschütz et al., 2014). If the relevant information for detecting such changes is predominantly contained in these rapid neural responses, it will be crucial whether it arrives at a favorable or unfavorable phase. Arriving at an unfavorable phase would naturally lead to a larger neural response latency which could delay successful change detection. Indeed, it has been found that larger response latencies are strongly correlated with longer reaction times, possibly caused by such an effect (Galashan et al., 2013). Evidence for a critical time window was also given by Ni et al., 2016 who demonstrated that both neural responses and reaction times were modulated by the gamma phase at which a sudden stimulus change occurred, and also depended on the V1-V4 interareal coherence (Rohenkohl et al., 2018).

Taken together, our findings directly demonstrate that signals carrying information of attended stimuli occur in short packages, tightly locked to the phase of the gamma-band oscillation, in the vicinity of the excitation maximum of the local target population. The results strongly support previous evidence for differential phase synchronization as a mechanism for attention-dependent selective signal routing. In particular, we established the methods to infer and quantify the properties of pulsed transfer schemes in neural data. These techniques will allow future studies to pinpoint similar processes in other visual areas, and to investigate whether the dynamical features exhibited by our data point towards a general principle for flexible information processing throughout the brain.

## Methods

### Experimental model and subject details

All procedures and animal care were in accordance with the regulation for the welfare of experimental animals issued by the federal government of Germany and were approved by the local authorities. Data from two adult male rhesus monkeys (Macaca mulatta) were used for this study. Parts of the data have been used in a previous publication (Grothe et al., 2018).

### Surgical procedures and behavioral task

Details about the surgical preparation, behavioral task and recording have been reported previously (Grothe et al., 2012). In short, animals were implanted under aseptic conditions with a post to fix the head and a recording chamber placed over area V4. Before chamber implantation, the monkeys had been trained on a demanding shape-tracking task (Fig. 2A). Neural signals were measured from area V4 with 1–3 epoxy-insulated tungsten microelectrodes (125 μm diameter; 1–3 MΩ at 1 kHz; Frederic Haer). Reference and ground electrodes for Monkey F were platinum-iridium wires below the skull at frontal and lateral sites. The reference for Monkey B was a platinum-iridium wire placed posteriorly below the skull, and the ground was a titanium pin at the posterior end of the skull.

The task (Fig. 2A) required fixation throughout the trial within a fixation window (diameter 1–1.5 degrees of visual angle) around a fixation point in the middle of the screen. Eye position was monitored using a video based eye tracking system (monkey B: custom made, monkey F: IScan Inc). After a baseline period, the monkeys had to covertly attend to one of two statically presented, closely spaced stimuli (shapes) that was cued (static/cue period). Then both shapes started morphing into other shapes. The monkeys were trained to respond by releasing a lever when the cued initial shape reappeared at a pseudorandomly selected position in the shape sequence, after two to five morph cycles. The animals had to ignore reappearance of the initial shape in the distracter sequence. The shapes were placed at equal eccentricity. Stimuli were presented with a refresh rate of 100 Hz on a 22 inch CRT monitor containing 1152 × 864 pixels (monkey B) or 1024 × 768 pixels (monkey F), which was placed at a distance of 92 cm (monkey B) or 87 cm (monkey F) in front of the animal.

In order to be able to track the information content of the stimuli within the neural activity, we used filled shapes and tagged the neural activity they evoke with imposed broadband luminance fluctuations (’flicker’) on the stimuli: we changed the luminance of the shapes by choosing a random, integer gray pixel value with each frame update of the display. For monkey F, the values were drawn from an interval [128, 172], and for monkey B, from the full range [0, 255], corresponding to luminance fluctuations in a range of 6.9-12.5 and 0.02-38.0 Cd/m^2^, respectively. Both shape streams had their own independent flicker time series of luminance values. Note that the flickering of the stimuli was not relevant to perform the task. A few trials were included in which only one stimulus was presented for offline controlling of response strength to individual stimuli.

### Recording

The electrodes’ signals were amplified by a factor of 1000 (Monkey B) or 5000 (Monkey F; MPA32I and PGA 64, 1-5000 Hz, Multi-Channel Systems GmbH) and digitized at 25 kHz (Monkey B: USB-ME-256 System, Multi-Channel Systems GmbH, Monkey F: A/D converter board, Multichannel systems and C++ based custom-made data acquisition system). For positioning stimuli, receptive fields were mapped manually while the monkey was fixating centrally, followed by an automated mapping procedure consisting of rapid presentations of circular dots. Stimuli were placed at equal eccentricity in the RF such that they induced similarly strong gamma-rhythmic activity (for details see Grothe et al., 2018). This requirement was successfully fulfilled in 16 recording sessions, resulting in 35 recording sites in total (Monkey F: 23 sites, Monkey B: 12 sites).

### Data preprocessing and site selection

From the recorded raw data, we extracted the local field potential (LFP) as a proxy reflecting average neural activity around the recording site, and the entire spiking activity (ESA) as a measure for more local spiking activity. All filters used in the process were realized as forward-backward FIR-filters to preserve the phase of the original signal, using the function ‘eegfilt’ from EEGLab (Delorme et al., 2011) with standard parameters for the cutoff:

- For obtaining the LFP signal, we extracted the low frequency component from the 25 kHz raw neural recordings by applying a bandpass FIR filter with band stops at 1 and 200 Hz.
- For obtaining the ESA signal, the raw data was first bandpass-filtered from 400 Hz to 2500 Hz. We then took the absolute value of the result, and subsequently applied a second bandpass FIR filter with band stops at 1 and 200 Hz to simplify phase dissociation (see detailed explanation below).

Finally, both signals were down-sampled to 1000 Hz and z-score normalized, yielding *y*_*LFP*_ and *y*_*MUA*_ used in our data analysis explained in the next section.

For ensuring that the recording sites were within the superficial layers of V4 we employed additional selection criteria:

- The shape of the visual evoked potential (VEP) caused by the stimulus onset shows the characteristic time course expected for the superficial layers, which starts with a negative deflection as opposed to the initial positive deflection observed in the deeper layers (for details see Givre et al., 1994; Nandy et al., 2017). We computed the VEP for each recorded site by averaging its LFP over trials.
- ESA and LFP exhibit spike-field coupling in the gamma frequency range. Spike-field coherence in the gamma frequency range has been reported to be high in the superficial layers and absent in the deep layers (Buffalo et al., 2011). For our analysis, spike-field couping was necessary in order to assess the ESA-aligned phase as a proxy for excitability’s phase.

These selection criteria left 11 recording sites for monkey F and 9 for Monkey B.

The experimental setup involved having two stimuli within the V4’s RF; each location was cued to be attended for half of the recorded trials in each session. We decided to split the trials of each recording site by the attended location, providing us with 22 data sets for monkey F and 18 data sets for monkey B. This was possible because 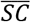 values from trial sets split by attended location were as statistically independent as 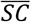 values from different recording sites, suggested by the fact that the distributions of the differences between the split sets and the full sets were not significantly different (Wilcoxon signed rank test).

### Data analysis

Data analysis consisted of several processing steps applied to the LFP and ESA signals in sequence. Briefly, first gamma phase and amplitude were extracted from the LFP, and gamma phase converted to excitability phase by assessing the spike-field coupling from ESA and LFP activity. Then, gamma excitability phase was used to isolate the signal values occurring at specific phases to form a phase-specific signal. Subsequently, periods of high (low) gamma amplitude were selected to form an amplitude-specific signal. In the final step of the analysis, stimulus information content in the neural signals was assessed by computing the spectral coherence between the luminance flicker and the dissociated LFP or ESA signals. In the following, each of those steps are explained in detail.

#### a) Extraction of gamma phase and amplitude

By applying a wavelet transform with Morlet kernels, we first computed the average power spectrum of the LFP signal *y*_*LFP*_ during the period in which the stimuli were morphing until 200 ms before a correct response, and normalized it by the average power spectrum observed before stimulus onset in the baseline period (separately for each recorded site). The spectra revealed clear peaks in the gamma frequency range, of which we extracted a lower and upper frequency limit by taking location at half of the highest point around the peak (approximately 40 to 100 Hz for monkey F recordings, and 50 to 110 Hz for monkey B). Subsequently, gamma activity was obtained by applying a forward-backward FIR band-pass filter to the LFP with cutoff frequencies determined by the lower/upper frequency limits. By applying the Hilbert transform to the result, we obtained LFP’s gamma phase Φ_*LFP*_ and amplitude A_*LFP*_.

Next, we would like to know when in each gamma oscillatory cycle neuronal activity is maximal, as a proxy for maximal excitability. Ideally, phases of high (or low) excitability should roughly correspond to high (or low) spiking activity. Unfortunately, the recorded ESA reflects only a small number of neurons next to the recording site, resulting in a signal that is too noisy to reliably extract gamma phase and amplitude. On the other hand, while the LFP provides a clean and reliable measure of the local populations’ rhythmic activity, its recording is affected by conduction delays and phase-shifts that depend on the recording electrode impendence as well as its precise location and orientation within the neural tissue (Bédard et al., 2004; Bédard and Destexhe, 2012; Gabriel et al., 1996; Nelson et al., 2008), making it a poor proxy for excitability. To resolve this issue, we related gamma phase information from the LFPs to spiking activity contained in the ESA by computing the mean ESA value for each LFP gamma phase, thus obtaining an estimate for the spike-field coupling. By subtracting from Φ_*LFP*_ the phase for which spike-field coupling was maximal we acquired the ESA-aligned gamma phase Φ_*ESA*_, which served as a proxy for excitability gamma phase throughout the entire analysis. For the amplitude of excitability’s gamma rhythm, we kept LFP’s amplitude A_*LFP*_.

#### b) Phase and amplitude dissociation, wavelet transform

Extraction of gamma phase-specific components of the neural recording signals *y* was performed before using a wavelet transform *W*_*f*_ to obtain a frequency-resolved neural signal representation, while dissection with respect to gamma amplitude was performed thereafter. Applying these three operations in sequence yields the dissected and spectrally resolved neural activity 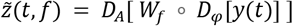. Here we write amplitude and phase dissection as formal operations *D*_*A*_ and *D*_φ_, respectively, which were realized as follows.

For performing phase dissociation, we first determined the time points *t*_*kφ*_ at which the excitability phase passed through a desired target phase (e.g., *φ* = 0 for peaks, or *π* for troughs). These times were then used to create a new signal *D*_*φ*_[*y*] by sampling from the original signal *y*(*t*) (LFP or ESA) at those points. In conjunction with the then following integration over time during the wavelet transform, we can formally write this notching operation by using the *δ*-distribution:

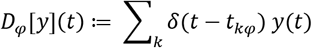

Note that in order to obtain the phase-dependency curves of the SIC, we evaluated it for a finite set of gamma phases with equal spacing. For this reason we wanted the ‘notched’ signal *y*(*t*_*kφ*_) to represent not only activity at exactly the time point *t*_*kφ*_, but also in its vicinity. This was trivially the case for the LFP since it was originally obtained by low-pass filtering. For the more rapidly varying ESA, we had to apply a second bandpass filter (see description above), in order to avoid ‘missing’ an activation peak by notching the signal at a slightly different time.

Amplitude dissociation was realized by first obtaining the distribution of gamma amplitudes throughout each individual recording session. From this distribution, we selected the 70th (and 30th) percentiles to use as high (and low) amplitude thresholds *A*_*hi*_ (and *A*_*low*_). Using these thresholds, we selected time periods exhibiting high (or low) oscillation amplitudes by means of indicator functions

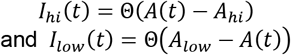

where Θ denotes the Heaviside function. Using these indicator functions, the amplitude specific spectra *D*_*A*_[*ỹ*](*t*, *f*) takes the form

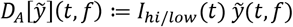

If only phase dissection was performed (no amplitude selection), we used the identity function for *D*_*A*_, thus *D*_*A*_[*ỹ*] = *ỹ*, and if only amplitude dissection was performed (no phase selection), we used the identity function for *D*_*φ*_, thus *D*_*φ*_[*y*] = *y*.

#### c) Spectral Coherence

To evaluate how much the luminance fluctuation signal *x*(*t*) of a shape contributed to the neural activity z(*t*), we utilized spectral coherence (SC). First, we computed the spectrograms 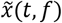 and 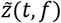 where *f* is the frequency and *t* is the time, using a wavelet transform with Morlet kernels. Here 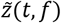 represents the neural signals which already underwent phase- and/or amplitude dissociation in conjunction with the wavelet transform as described in the preceding section. The transform yields complex valued coefficients representing the amplitude and phase of the signals. By evaluating the normalized cross-correlation between 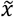 and 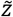 we obtained the spectral coherence measure

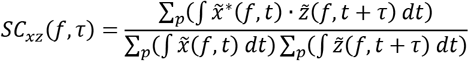

where *τ* is the lag between the two signals, and where the sums are performed over the population of trials *p* included in the computation.

Due to the normalization terms in the denominator, the values of *SC*_*xz*_ lie between zero and one. All integrals were computed over all times for which *t* and *t* + *τ* lie within a selected time period during a trial, i.e. from the beginning of the second morph cycle until 200ms before the monkey’s response. Summation was performed either over trial repetitions (*r*) and recording sites (*s*) for population analyses, *p* = {*r*, *s*}, or over trial repetitions *only* for analysis of *single* sites, *p* = {*r*}.

Once *SC* is calculated, we compute the pooled value 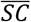 over a region of interest in frequency-time lag space to reduce a two-dimensional result to a single value. The region of interest was defined as a frequency-dependent cone of width 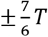 around 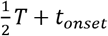, where *T* = 1/*f* and *t*_*onset*_ denotes the onset delay of the neural response in V4 after stimulus onset which was 50ms in monkey F and 60ms in monkey B (see Grothe et al., 2018). We first took the average across lags within the frequency-dependent region of interest, and then took the mean of the time averages from 5 up to 15 Hz. 15 Hz was selected as upper the limit, since the majority of the individual sets results did not yield significant *SC* above this value.

#### d) Confidence intervals and statistical tests

For assessing significance of each 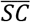 measure, we computed the 95% chance level of its value being different from zero (indicated by the gray shading towards the bottom of each plot in figures 4A, 5A and 6). This was done by taking the 95th-percentile from the distribution of 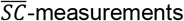 gathered by pairing the neural recording signal with 200 surrogate luminance flicker signals.

95% confidence intervals for 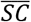 were gathered via bootstrapping. 20000 new sets of trials were generated by resampling with replacement from the available trials, each resampled set containing the same number of trials as the original (resulting intervals are indicated by shading around the 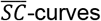 in figures 4A and 6, and by error bars in figure 5A).

For assessing whether two individual 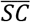 measurements are significantly different, i.e. for all the cumulative results in figures 4A, 5A and 6 as well as for each individual set in the scatter plots of figures 4B and 5B, a non-parametric permutation statistical test was utilized: we constructed the null distribution for 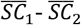 with 20000 repetitions, resampling from the available trials without replacement (Maris et al., 2007; Maris and Oostenveld, 2007), and checked whether the probability to obtain a value larger than the true difference was less than 5%.

To assess whether the distributions of 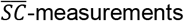 for individual data sets are significantly different between two conditions (i.e. whether the data clouds in the scatter plots in figures 4B and 5B lie above or below the diagonal), we determined whether the ratios between the 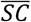 values for individual sets are significantly different from 1 by using the Wilcoxon signed rank test on the distribution computed via 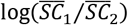.

## Author Contributions

Conceptualization, D.L. and U.E.; Methodology, D.L.; Software, D.L.; Formal Analysis, D.L.; Investigation, I.G., A.K.; Writing – Original Draft, D.L. and U.E.; Writing – Review & Editing, D.L., I.G., A.K., U.E.; Visualization, D.L.; Supervision, U.E. and A.K.; and Funding Acquisition, A.K. and U.E.

## Acknowledgements

The authors thank K. Thoß, R. Hakizimana, and K. Taylor for monkey care and training; D. Rotermund for support in data preprocessing. This work was supported by the BMBF (Bernstein Group for Computational Neuroscience Bremen, Grant 01GQ0705, Innovationswettbewerb Medizintechnik, Grant 01 EZ 0867, Bernstein Award Udo Ernst, Grant 01GQ1106), the DFG (Priority Program 1665, Grant ER 324/3 and KR 1844/2-2), the University of Bremen’s Research-Focus Neurotechnology, Creative Unit I-See, and Zentrum fuer Kognitionswissenschaften, and the Leibniz Graduate School for Primate Neurobiology (to I.G.).

